# Statistical analysis and simulation allowing simultaneously positive, negative, and no crossover interference in multilocus recombination data

**DOI:** 10.1101/2022.11.02.514815

**Authors:** Shaul Sapielkin, Zeev Frenkel, Eyal Privman, Abraham B. Korol

**Affiliations:** The Department of Evolutionary and Environmental Biology, University of Haifa, Haifa, 3498838, Israel; Institute of Evolution, University of Haifa, Haifa, 3498838, Israel

**Keywords:** Crossover interference, coefficient of coincidence, gamma model

## Abstract

Crossover interference (COI) is a widespread feature of homologous meiotic recombination. It can be quantified by the classical coefficient of coincidence (CoC) `but this characteristic is highly variable and specific to the pair of chromosomal intervals considered. Several models were proposed to characterize COI on a chromosome-wise level. In the gamma model, the strength of interference is characterized by a shape parameter *ν*, while the gamma-sprinkled two-pathway model (GS) accounts for both interference-dependent and independent crossover (CO) events by fitting a mixture of gamma distributions with *v*>1 and *v*=1, correspondingly, and mixture proportions 1-*p* and *p*. In reality, COI can vary along chromosomes resulting in low compliance of the fitted model to real data. Additional inconsistency can be caused by common neglecting of possible negative COI in the model, earlier reported for several organisms. In this work, we propose an extension of the GS-model to take possible negative COI into account. We propose a way for data simulation and parameter estimation for such situations.

## 1 Introduction

Crossover interference (COI) is the deviation of the double crossover (CO) rate from the expected frequency assuming the independence of CO events (chiasmata) along the chromosome during meiosis. The phenomenon was discovered in the early experiments on recombination in *Drosophila melanogaster* [26, 35, 43]. The degree of COI can be quantified by the coefficient of coincidence (CoC), the ratio of the observed to the expected number of double COs for a pair of chromosomal intervals: *CoC* < 1, *CoC* > 1, and *CoC* = 1 corresponding to positive, negative, and no COI, respectively. The nature of COI was explained in two ways: (i) as the result of mutual interaction, either repulsion or clustering, of COs (referred to as chiasmata interference) [36], and (ii) as non-random involvement of chromatids in neighboring CO exchanges (referred to as chromatid interference) [26]. Studies in many species [11, 13, 27, 31, 32, 44, 47] showed no significant effect of chromatid interference on COI, so most of the proposed COI models assume no chromatid interference [21, 22, 33, 47].

To describe interference as a single mechanism of CO formation acting along the chromosome or at the entire genome level, one of the most conveniently applicable is the gamma model [5, 10, 19, 21, 48]. The gamma model is a common method to describe interference as a single mechanism of CO formation acting along the chromosome or at the level of the entire genome. It considers the dependence of the CO number distribution per individual, or the distances between COs, on the interference “strength” parameter (*v*). The values *v*>1 correspond to positive COI (“repulsion” between COs), while *v*=1 implies no interference. The gamma model was further modified corresponding to the discovery of the interference-independent metabolic pathway of meiotic recombination alongside the common interference-dependent pathway [24]. Thus, the resulting “gamma-sprinkled” (GS) model assumes a mixture of gamma distributions with *v*>1 and *v*=1 (cases of *positive* COI and *no* COI) with proportions 1-*p* and *p*, respectively, and appeared to be more consistent with empirical data [38]. Yet, cases of possible negative COI were also reported on different organisms (*Neurospora*, yeast, *Drosophila, Arabidopsis*, maize, barley, wheat), and the possible presence of CO clustering effect (negative COI) was reported [1, 2, 39, 42, 45, 3, 4, 8, 9, 12, 16–18] When applied to such cases, the GS-model gives biased estimates of both parameters, *p*, and *v*, due to the presence of three components (with *v*>1, *v*=1, and *v*<1). Therefore, to take account of possible negative interference, we extend the GS model by fitting a mixture of the corresponding three distributions.

## 2 Statistical analysis of COI landscape using *CoC* calculations on high-density segregation data

In general, for a pair of adjacent intervals on a genetic map flanked by three points (genetic markers, m1-m2-m3), the log-likelihood function for a testcross progeny can be presented as:

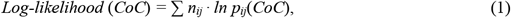

where *p*_*ij*_ are the probabilities of gametes: *p*_00_ = (1*-r*_1_*-r*_2_*+CoC*·*r*_1_·*r*_2_) for no COs on both intervals, *p*_10_ = (*r*_1_*-CoC*·*r*_1_·*r*_2_) and *p*_01_ = (*r*_2_*-CoC*·*r*_1_·*r*_2_) for CO on the 1^st^ and the 2^nd^ intervals, respectively and *p*_11_ = (*CoC*·*r*_1_·*r*_2_) for double COs; while 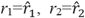 are corresponding recombination fractions and *n*_ij_ are the observed number of cases for the corresponding classes and calculated separately for each interval independently.

With a relatively small sample size, using too short intervals implies insufficient statistical power of discriminating the alternatives H_1_(*CoC*≠1) versus H_0_(*CoC*=1). Yet, in such situations, increasing the interval size by using only a subset of markers may lead to a classification problem for each genotype as “CO” or “non-CO” for these larger-length intervals, because of possible double COs that will be classified as “non-CO”. A slight modification of the definition “CO genotype” for each new interval solves this problem. Namely, we define a recombinant genotype in a larger-length interval as any genotype where one or more CO events have occurred in the corresponding sub-intervals contained within the larger interval (when considering all of the markers). We refer to this type of classification as the “crossover superposition principle”, or CSP.

## 3 Heterogeneity COI landscape by local *CoC* calculations from high-density segregation data

In our estimation of *CoC*, we maximize the likelihood function (EM algorithm) [15] based on the Nelder-Mead optimization (simplex method) [37, 46] for three hypothetical situations, where recombination occurs: (1) without COI according to Haldane Mapping Function, i.e., *CoC* =1 [25]; (2) with COI according to the Kosambi Mapping Function, i.e. *CoC* =2*r* [30]; and (3) with arbitrary *CoC*. As a real data example, we employ dataset Cluster #3 (842 genotypes) of the Norway spruce (*Picea abies*) high-density segregation data published by Bernhardsson and co-authors [6]. For the analysis of interference, we constructed high-quality maps using MultiPoint-ultradense software [40]. To classify the genotypes into 0-0, 0-1, 1-0, and 1-1 groups we used the approach described in the previous section with different sizes of interval pairs window (*l*_*1*_ +*l*_*2*_) along linkage groups from 10 to 70cM. The obtained vectors of frequencies *P*= (*p*_00_, *p*_01_, *p*_10_, *p*_11_)=*P*(*l*_1_,*l*_2_) were employed for local maximum likelihood (ML) estimation of *CoC* with sliding windows along the linkage map and likelihood-ratio (LR) test to COI models compliance. On LG7 we found a region with the relaxation of positive COI followed by no-COI and even negative COI at some map positions (Fig.1).

**Fig. 1.**
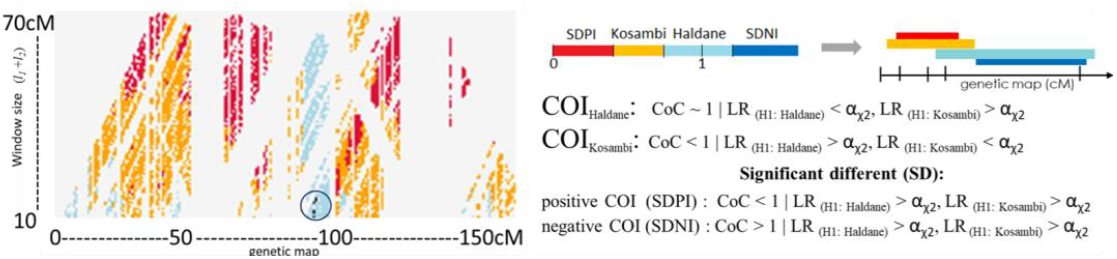
COI landscape along the genetic map of Norway spruce LG7 (data from [6]) with several sizes of intervals and compliance of COI to hypothesis tests. COI: positive, significantly stronger than Kosambi – red; Kosambi – orange; Haldane – blue; significant negative COI - dark blue (additionally marked with a circle).

To analyze COI at the chromosome-wide level with the GS-model on the same data, we used CODA software [23] simulations to estimate the GS model parameters (Fig 2). The solution obtained based on simulations points to the presence of a non-interfering component in the distribution of recombination events, with a proportion *p* = 0.29. Likewise, the real data show an excess of gametes with CO > 3 (Fig 2, right) and an excess of short inter-crossover (CO-CO) distances (Fig 2, left). The observed deviation of the CODA simulations using the obtained estimates of *v* and *p* parameters, i.e., excess of both multiple exchanges and short inter-CO intervals, remarkably coincides with our finding of possible negative COI for the same linkage group (see Fig 1). However, a more objective conclusion on how to interpret the mentioned deviations from the GS-model requires using an extended GS-model allowing for the possibility of negative interference.

**Fig. 2.**
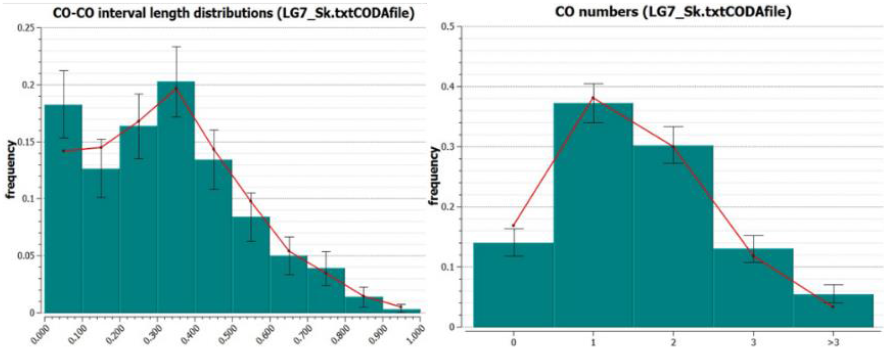
CODA simulations using GS-model for LG 7 Norway spruce data from [6]. Left - CO-CO intervals length distribution; right - CO number per individual distribution. Obtained estimates of GS parameters: *v*= 7.90, p= 0.29; red – simulation, green – real data.

## 4 Extension of the GS-model to take possible negative COI into account

### 4.1 Genetic COI model based on the gamma distribution

Let genetic recombination events arise consequently on the chromosome with distance from the previous recombination event depending only on the position of this previous event. Let distances *D* on the genetic map between recombination events follow a gamma distribution with the parameters *v* (shape) and *θ* (scale) [48]:

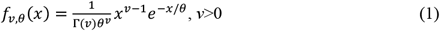

This model can be considered as a generalization of the classical Haldane model, where CO events are independent, hence the distances between consequent events obey exponential distribution:

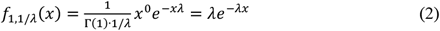

From Γ(1,1/λ)∼*Exp*(λ) follows the Haldane model. If 2*v* is an integer number, then gamma distribution coincides with χ^2^ distribution with 2*v* degrees of freedom (*v*=*df*/2, *θ*=2):

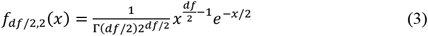

With *df*=4 and *v*=2, *θ* =100/2, the gamma distribution coincides with the χ^2^ distribution with *df*=2. Such distribution of distances between COs is close to the Kosambi model [30].

In general, the average and variance of the distances between consequent COs fitting the gamma model can be presented as:

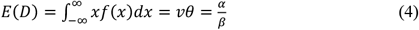

and

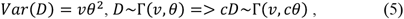

where *c* is a scaling coefficient.

Genetic distance (in Morgans) is the expected number of CO events between two points on the genetic map. In terms of cM distances: *E*(*D*)=100=*vθ*. Hence, *θ*=100/*v* and to specify the model we need only parameter *v*. The case with *v*=1 corresponds to the situation of no COI, *v*>1 implies positive COI, and 0<*v*<1 - negative COI.

### 4.2 Simulation of distances between two CO events based on gamma distribution with parameter *v*

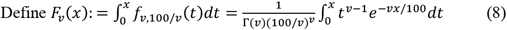

Let *u* be a random variable with uniform distribution on the interval (0, 1): *U*(0,1).

We simulate *D*_*v*_ as 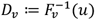

For calculations of *Fv*(*x*) we use the following approximation:

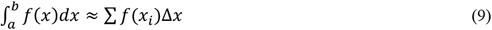

Note that

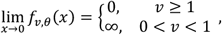

therefore, such an approximation is problematic for *v*<1 and small *x*_*i*_. For such cases, we can use the following asymptotic:

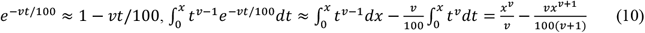

According to expressions 8-10, we used the following model for CO events. Let the process be independent of the position and orientation of the simulated region on the genetic map. Based on the symmetry, the distribution of distances from the start of the genetic map to the first CO event is as in the Haldane model. We simulate positions *x* of CO events on the map [0, *L*] (*L* is the length of the map scored in cM) as the following:

Let *u*^(*i*)^∼U[0,1],

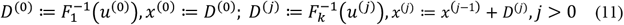

We stop the simulation when *x*^(*j*+1)^ > *L*. Only values *x*^(*n*)^ from [0, *L*] are considered as positions of CO events on the chromosome. Distance *D*^(0)^ is simulated according to the Haldane model because of an absence of the previous CO event (hence, no dependence like in the Haldane model). The resulting ordered set of values *x*^(*i*)^ (vector) is denoted by *X*.

### 4.3 Simulation of CO events based on a mixture of three processes (with parameters *v*_0_, *v*_1_, and *v*_2_)

Let there are three types of CO events: dependent according to the models with parameters *v*_0_, *v*_1,_ and *v*_2_, and events of one type are independent of the events of the other types. Let *p*_1_ and *p*_2_ be the proportion of events of type 1 and 2 on the chromosome, 0 ≤ *p*_1_, *p*_2_ ≤ 1, *p*_1_ + *p*_2_ ≤ 1. If 0<*p*_1_,*p*_2_,*p*_1_*+p*_2_<1, then to maintain on average 0.01 CO rate per 1 cM, the following condition should hold: *p*_1_*ν*_1_*θ*_1_ + *p*_2_*ν*_2_*θ*_2_ + (1 − *p*_1_ − *p*_2_)*ν*_0_*θ*_0_ = 100, so that

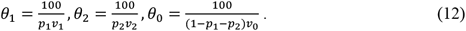

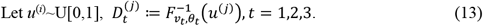

Our simulations are based on iterations of 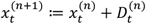. Simulations continue while 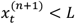, i.e. only values 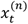 from [0, *L*] are considered as positions of COs on the chromosome. We unite three sets of the obtained values and sort them by increasing. The resulting vector is denoted by *X*.

### 4.4 Simulation of CO events with changing parameter *v* along the chromosome

Let parameter *v* vary along the genetic map: *v*=*v*(*x*). Define 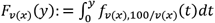. The distribution density of the distance to the next CO event (after the CO event at *x*_0_) is 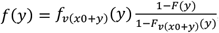, where 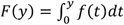. Hence, 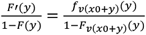 and 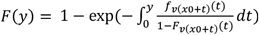. Analogously to the case with constant *v*, we simulate positions of CO events on the map [0, *L*] as the following:

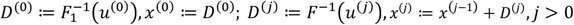

where *u*^(*i*)^∼U[0,1].

As an illustration of an application of the described approach, we generated a segregating test-cross data set with characteristics similar to Norway spruce real data mentioned above (see section 3). Several population sizes were simulated (*n*=200, 500, 1000, and 5000); with chromosome zones of varying *v*: strong positive COI (*v*=5), Kosambi (*v*=2), Haldane (*v*=1), negative COI (*v*=0.5) and strong negative COI (*v*=0.2). We analyzed the distribution of the *CoC* using two approaches: CSP (described in section 3) and standard (based on flank markers). For population sizes *N*= 200 and 500 we employed window sizes *l*_*1*_ +*l*_*2*_ = 20 cM and 30 cM, while 10 cM and 20 cM window sizes were employed for *N*=1000 and 5000, bearing in mind the increase in resolution with a higher sample size (Fig 3). We found that for cases of strong negative COI, *CoC* estimates based on CSP genotypes classification are more accurate than those obtained from the standard classification.

**Fig. 3.**
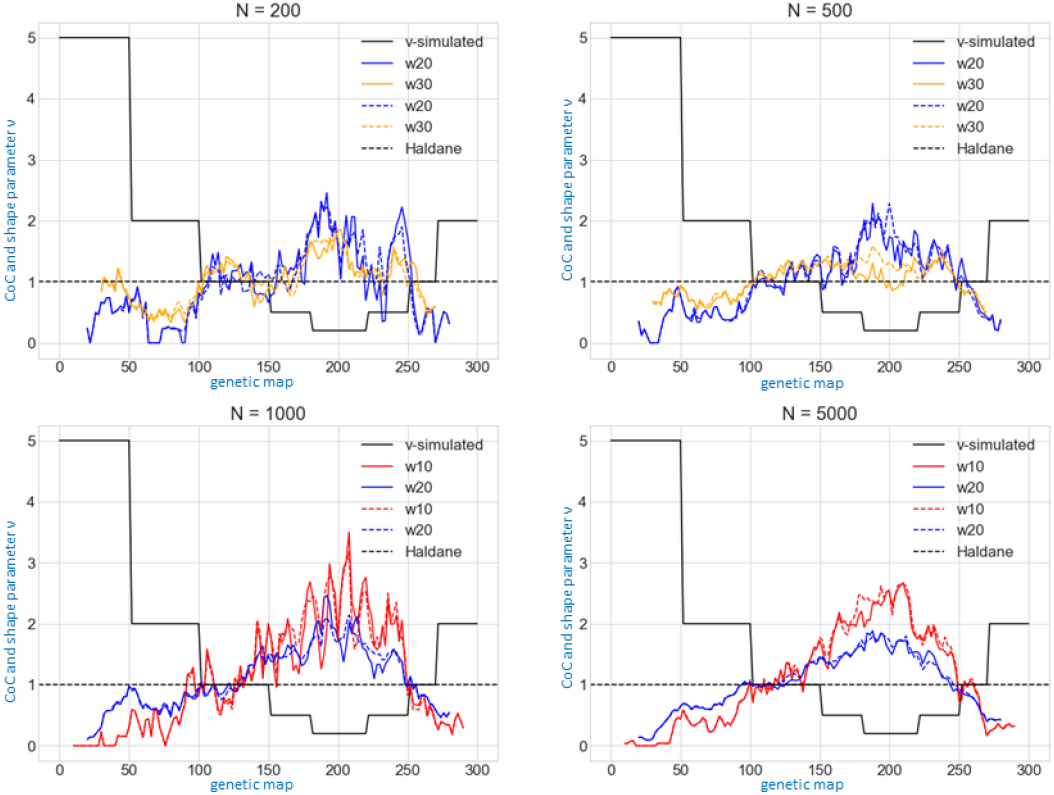
Simulation of COI landscape with variable *v*. Simulated *v* is represented by step blackline, variation in *CoC* estimates based on flanked-markers and CSP genotype classification are shown by solid and dotted lines, respectively, for each size (w=10, 20, 30 cM) of the sliding window (see section 4).

## 5 Statistical analysis of COI using the modified GC model

### 5.1 Basic statistics for a single vector and a set of vectors of CO positions

Let *X* be a vector of consequent CO positions on a chromosome for an individual of a segregating population. The dimension of this vector (dim(*X*)≥0) corresponds to the number of CO events for the individual. Let further denote the set of lengths between consequent CO events by

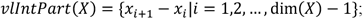

if dim(*X*)≤1 then the set *νlIntPart*(*X*) is empty. Let ***X*** be the set of vectors *X* for the entire population. Then we denote by *N*_*m*_(*X*) the number of vectors *X* from set ***X*** such that dim(*X*) = *m*, so that 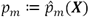 is the proportion of vectors *X* with dimension *m* and ***p*** = ***p***(*X*) = (*p*_0_, *p*_1_, …) is a vector of observed proportions of individuals with different numbers of CO; in practice, we unite all cases with *m* ≥ 5. We denote the observed cumulative distribution function for values from ⋃_*X*∈*X*_ *νlIntPart*(*X*) by *F*(*x*)*=F*(*x*, ***X***).

### 5.2 Distribution of the considered statistics

For a given chromosome length *L* and parameters *v*_0_, *v*_1_, *v*_2_, *p*_1_, *p*_2_ we simulate a large enough (e.g., *N*=5000) set *X*_0_ = *X*_0_(*L*, *ν*_0_, *ν*_1_, *ν*_2_, *p*_1_, *p*_2_, *N*) of vectors *X*. Based on this set we generated the “expected” distributions for the H_0_ hypothesis:

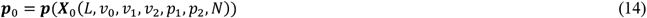

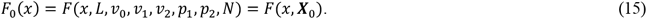

### 5.3 Testing the model adequacy

For the map positions of all CO events observed on *n* individuals, we can define the vector *X*_*obs*_ = (*X*_1_, …, *X*_*n*_) and compare two alternative hypotheses for the observed results:

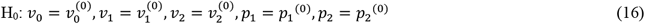

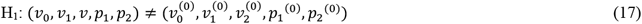

In particular, we are interested to compare H_0_ (no COI) versus various alternatives, like H1 only positive and only negative COI, positive + neutral, negative + neutral, and a mixture of all three cases) We employed two statistical tests for comparing such alternatives, the χ^2^ test based on CO number and the Kolmogorov–Smirnov test based on the cumulative distribution of distances between CO events.

χ^2^ test based on CO number: Under H_0_, the test statistics

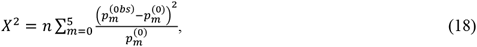

should follow 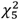 distribution. Thus, the p-value for the test is defined by 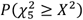.

By Kolmogorov–Smirnov test:

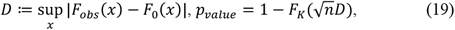

where *F*_*K*_(*x*) is the cumulative distribution function for the Kolmogorov distribution

### 5.4 Searching for a confident set of parameters

The motivation of our approach is to estimate the contribution of three putative mechanisms of CO localization (COI-free, positive COI, and negative COI) to the observed patterns of multilocus recombination in real data. We assume, that the three mechanisms may work simultaneously and we model such situations by the mixture of three processes (section 4), with parameters *v*_0_=1 (COI-free), 0<*v*_1_<1 (COI negative), and *v*_2_>1 (COI positive). For various parameter combinations (*v*_0_, *v*_1_, *v*_2_, *p*_1_, *p*_2_), we calculate *p*-values for both methods (section 5.3). Only model(s) with not too small *p*-values for both methods should be considered as a relevant description of the considered data. Such a method enables estimating model parameters assuming homogeneity of the entire process along the chromosome or large enough chromosomal segments. This may not be the case for real data (see example in Fig.1). Yet, the following simulated example shows that this approach can generate reasonable results. We simulated a testcross progeny with 2000 genotypes and 200 markers on a chromosome of 200 cM length. In the simulations, the following parameter values were employed: *v*_0_=1, *v*_1_=0.5, *v*_2_=2, *p*_1_=0.2, *p*_2_=0.4, *p*_0_=1-*p*_1_-*p*_2_=0.4. Using the described procedure (5.1-5.3), the best set of parameter estimates was: 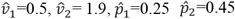. We tested the hypothesis H_1_ that parameter values are equal to the estimated ones against H_0_ that *p*_1_=0, *p*_2_=1 (i.e., assuming unimodal distribution with the presence of only positive COI). Under 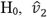 was equal to 1.10. The distribution of CO numbers in our simulated data significantly (*p*-value<0.001) differed from the one expected under H_0_. Therefore, the proposed analysis is able to detect the mixed nature of interference patterns despite the aforementioned simplifying assumption of homogeneity.

## 6 Conclusions and prospects

Crossover interference along a genetic map can be investigated at local and chromosome-wide levels. Both these approaches have limitations. In many practical situations, COI can vary along chromosomes resulting in low compliance of the chromosome-wide model to real data. For a qualitative assessment of the local COI, the standard analysis using CoC highly depends on the sample size and the size of the pair of intervals under study. To address the common problem of insufficient sample size for COI detection, we proposed to employ larger size intervals for CoC analysis. With this approach, we consider a genotype as recombinant in a larger-length interval if one or more CO events have occurred in the sub-intervals contained within the larger interval. We demonstrated that, as expected, CoC estimates for this pragmatic classification of genotypes are more accurate compared to those resulting from the standard classification. This makes it possible to better assess the impact of the negative COI phenomenon on the local area of the genetic map. To check for the robustness of the resulting conclusions, we perform such analysis on a range of sizes of the consequent intervals along a chromosome. Yet, bearing in mind the dependence of the CoC on the interval sizes for the same gamma model COI, a better approach would be to use the ML estimate of the corresponding point-specific gamma distribution parameter *v* (not shown, though we are developing this approach).

For the chromosome-wide COI, the currently widely employed GS-model does not allow testing for the presence of negative COI, which may exist as demonstrated by some empirical evidence. Due to the absence of such a possibility in the current versions of the GS-model, negative COI cases (if exist) are mixed with no-COI cases in the output results. In this paper, we proposed an extension of the GS-model aimed to take possible negative COI into account. For this purpose, we simulate crossover events as a mixture of three COI processes (positive, negative, and no COI). In this way, we estimate distributions of the CO number per genotype and inter-CO distances expected under this model. Comparisons of the distributions observed for real data with the massive simulations enable estimating the parameters of the model and testing the corresponding hypotheses. This approach will assist in the objective addressing of the question of the presence of negative COI for each suitable experimental data instead of ignoring it from the beginning. Usually, COI is considered as a factor preventing the formation of excessive COs [7]. Recent studies in Arabidopsis and some crops have shown that loss-of-function mutations in anti-recombination genes, that are involved in DNA repair of excessive double-strand breaks during meiosis, displayed a very high increase in the CO rates [14, 20, 34, 41]. Elucidating the possible role of de-regulation of COI in this phenomenon remains an open question. It is important to mention that in some earlier reports, including drosophila and wheat, NCOI was associated with pericentromeric regions [9, 16, 28, 29, 39] where normally recombination is strongly suppressed.

In the ongoing study on tetraploid wheat in our lab, the recombinogenic effect of a double mutation of the helicase gene *Recq4* in the A and B genomes (*recq4A-recq4B*) is tested. One of the questions is whether changes in recombination involve repatterning of COI including the possibility of NCOI [28].

## References

1. Aggarwal, D.D. et al.: Desiccation-induced changes in recombination rate and crossover interference in Drosophila melanogaster: evidence for fitness-dependent plasticity. Genetica. 147, 3–4, 291–302 (2019). https://doi.org/10.1007/s10709-019-00070-6.

2. Aggarwal, D.D. et al.: Experimental evolution of recombination and crossover interference in Drosophila caused by directional selection for stress-related traits. BMC Biol. 13, 1, (2015). https://doi.org/10.1186/s12915-015-0206-5.

3. Aggarwal, D.D. et al.: Seasonal changes in recombination characteristics in a natural population of Drosophila melanogaster. Heredity (Edinb). 127, 3, 278–287 (2021). https://doi.org/10.1038/s41437-021-00449-2.

4. Auger, D.L., Sheridan, W.F.: Negative crossover interference in maize translocation heterozygotes. Genetics. 159, 4, 1717–1726 (2001).

5. Bailey NTJ: Introduction to the mathematical theory of genetic linkage. The Clarendon Press, London (1961).

6. Bernhardsson, C. et al.: An Ultra-Dense haploid genetic map for evaluating the highly fragmented genome assembly of Norway spruce (Picea abies). G3 Genes, Genomes, Genet. 9, 5, 1623–1632 (2019). https://doi.org/10.1534/g3.118.200840.

7. Bomblies, K. et al.: The challenge of evolving stable polyploidy: could an increase in “crossover interference distance” play a central role? Chromosoma. 125, 2, 287–300 (2016). https://doi.org/10.1007/s00412-015-0571-4.

8. Bowring, F.J., Catcheside, D.E.A.: Evidence for negative interference: Clustering of crossovers close to the am locus in Neurospora crassa among am recombinants. Genetics. 152, 3, 965–969 (1999). https://doi.org/10.1093/genetics/152.3.965.

9. Boyko, E. et al.: A high-density cytogenetic map of the Aegilops tauschii genome incorporating retrotransposons and defense-related genes: insights into cereal chromosome structure and function. Plant Mol. Biol. 48, 5–6, 767–790 (2002). https://doi.org/10.1023/a:1014831511810.

10. Browning, S.: The relationship between count-location and stationary renewal models for the chiasma process. Genetics. 155, 4, 1955–1960 (2000). https://doi.org/10.1093/genetics/155.4.1955.

11. Chen, S.Y. et al.: Global Analysis of the Meiotic Crossover Landscape. Dev. Cell. 15, 3, 401–415 (2008). https://doi.org/10.1016/j.devcel.2008.07.006.

12. Colombo, P.C., Jones, G.H.: Chiasma interference is blind to centromeres. Heredity (Edinb). 79 (Pt 2), 2, 214–227 (1997). https://doi.org/10.1038/HDY.1997.145.

13. Copenhaver, G.P. et al.: Assaying genome-wide recombination and centromere functions with Arabidopsis tetrads. Proc. Natl. Acad. Sci. U. S. A. 95, 1, 247–252 (1998). https://doi.org/10.1073/pnas.95.1.247.

14. Crismani, W. et al.: FANCM limits meiotic crossovers. Science. 336, 6088, 1588–90 (2012). https://doi.org/10.1126/science.1220381.

15. Dempster, A.P. et al.: Maximum Likelihood from Incomplete Data Via the EM Algorithm. J. R. Stat. Soc. Ser. B. 39, 1, 1–22 (1977). https://doi.org/10.1111/j.2517-6161.1977.tb01600.x.

16. Denell, R.E., Keppy, D.O.: The nature of genetic recombination near the third chromosome centromere of Drosophila melanogaster. Genetics. 93, 1, 117–130 (1979).

17. Drouaud, J. et al.: Sex-specific crossover distributions and variations in interference level along Arabidopsis thaliana chromosome 4. PLoS Genet. 3, 6, 1096–1107 (2007). https://doi.org/10.1371/journal.pgen.0030106.

18. Esch, E., Weber, W.E.: Investigation of crossover interference in barley (Hordeum vulgare L.) using the coefficient of coincidence. Theor. Appl. Genet. 104, 5, 786–796 (2002). https://doi.org/10.1007/s00122-001-0842-8.

19. Falque, M. et al.: Two types of meiotic crossovers coexist in maize. Plant Cell. 21, 12, 3915–3925 (2009). https://doi.org/10.1105/tpc.109.071514.

20. Fernandes, J.B. et al.: Unleashing meiotic crossovers in hybrid plants. Proc. Natl. Acad. Sci. U. S. A. 115, 10, 2431–2436 (2018). https://doi.org/10.1073/pnas.1713078114.

21. Foss, E. et al.: Chiasma interference as a function of genetic distance. Genetics. 133, 3, 681–691 (1993). https://doi.org/10.1093/genetics/133.3.681.

22. Foss, E.J., Stahl, F.W.: A test of a counting model for chiasma interference. Genetics. 139, 3, 1201–1209 (1995). https://doi.org/10.1093/GENETICS/139.3.1201.

23. Gauthier, F. et al.: CODA (crossover distribution analyzer): Quantitative characterization of crossover position patterns along chromosomes. BMC Bioinformatics. 12, 1, 27 (2011). https://doi.org/10.1186/1471-2105-12-27.

24. Giraut, L. et al.: Genome-Wide Crossover Distribution in Arabidopsis thaliana Meiosis Reveals Sex-Specific Patterns along Chromosomes. PLoS Genet. 7, 11, e1002354 (2011). https://doi.org/10.1371/journal.pgen.1002354.

25. Haldane, J.B.S.: The combination of linkage values, and the calculation of distances between the loci of linked factors. J. Genet. 9, 299–309 (1919).

26. Haldane, J.B.S.: The Cytological Basis of Genetical Interference. Cytologia (Tokyo). 3, 1, 54–65 (1931). https://doi.org/10.1508/cytologia.3.54.

27. Hawthorne, D.C., Mortimer, R.K.: Chromosome Mapping in Saccharomyces: Centromere-Linked Genes. Genetics. 45, 8, 1085 (1960). https://doi.org/10.1093/GENETICS/45.8.1085.

28. Korol, A. et al.: Methods for Genetic Analysis in the Triticeae. Springer (2009). https://doi.org/10.1007/978-0-387-77489-3.

29. Korol AB, Preygel IA, P.S.: Recombination Variability and Evolution. London Chapman Hall. 361 pp (1994).

30. Kosambi, D.D.: The Estimation of Map Distances From Recombination Values. Ann. Eugen. 12, 1, 172–175 (1943). https://doi.org/10.1111/j.1469-1809.1943.tb02321.x.

31. Li, X. et al.: Dissecting meiotic recombination based on tetrad analysis by single-microspore sequencing in maize. Nat. Commun. 2015 61. 6, 1, 1–9 (2015). https://doi.org/10.1038/ncomms7648.

32. Lindegren, C.C., Lindegren, G.: Locally Specific Patterns of Chromatid and Chromosome Interference in Neurospora. Genetics. 27, 1, 1–24 (1942). https://doi.org/10.1093/GENETICS/27.1.1.

33. McPeek, M.S., Speed, T.P.: Modeling interference in genetic recombination. Genetics. 139, 2, 1031–1044 (1995). https://doi.org/10.1093/genetics/139.2.1031.

34. Mieulet, D. et al.: Unleashing meiotic crossovers in crops, (2018). https://doi.org/10.1038/s41477-018-0311-x.

35. Morgan, T.H. et al.: The genetics of Drosophila. Bibliogr. Genet. (1925).

36. Muller, H.J.: The Mechanism of Crossing-Over. Am. Nat. 50, 592, 193–221 (1916). https://doi.org/10.1086/279534.

37. Nelder, J.A., Mead, R.: A Simplex Method for Function Minimization. Comput. J. 7, 4, 308–313 (1965). https://doi.org/10.1093/comjnl/7.4.308.

38. Otto, S.P., Payseur, B.A.: Crossover Interference: Shedding Light on the Evolution of Recombination. Annu. Rev. Genet. 53, 19–44 (2019). https://doi.org/10.1146/annurev-genet-040119-093957.

39. Peng, J. et al.: Molecular Genetic Maps in Wild Emmer: genome-wide coverage, massive negative interference, and putative quasi-linkage. Genome Res. 10, 1509–1531 (2000). https://doi.org/10.1101/gr.150300.role.

40. Ronin, Y.I. et al.: Building ultra-high-density linkage maps based on efficient filtering of trustable markers. Genetics. 206, 3, 1285–1295 (2017). https://doi.org/10.1534/GENETICS.116.197491/-DC1.

41. Séguéla-Arnaud, M. et al.: Multiple mechanisms limit meiotic crossovers: TOP3α and two BLM homologs antagonize crossovers in parallel to FANCM. Proc. Natl. Acad. Sci. U. S. A. 112, 15, 4713–8 (2015). https://doi.org/10.1073/pnas.1423107112.

42. Sinclair, D.A.: Crossing over between closely linked markers spanning the centromere of chromosome 3 in Drosophila melanogaster. Genet. Res. 26, 2, 173–185 (1975). https://doi.org/10.1017/S0016672300015974.

43. Stevens, W.L.: The analysis of interference. J. Genet. 32, 1, 51–64 (1936). https://doi.org/10.1007/BF02982501.

44. Strickland, W.N.: An analysis of interference in Aspergillus nidulans. Proc. R. Soc. London. Ser. B - Biol. Sci. 149, 934, 82–101 (1958). https://doi.org/10.1098/RSPB.1958.0053.

45. Tsubouchi, T. et al.: The Meiosis-Specific Zip4 Protein Regulates Crossover Distribution by Promoting Synaptonemal Complex Formation Together with Zip2. Dev. Cell. 10, 6, 809–819 (2006). https://doi.org/10.1016/j.devcel.2006.04.003.

46. Wright, M.H.: Direct Search Methods: Once Scorned, Now Respectable. Pittman Res. Notes Math. Ser. Numer. Anal. 1995. 191–208 (1996). https://doi.org/10.2/JQUERY.MIN.JS.

47. Zhao, H. et al.: Statistical analysis of crossover interference using the chi-square model. Genetics. 139, 2, 1045–1056 (1995). https://doi.org/10.1093/genetics/139.2.1045.

48. Zhao, H., Speed, T.P.: On Genetic Map Functions. Genetics. 142, 4, 1369–1377 (1996). https://doi.org/10.1093/genetics/142.4.1369.

